# Degradation of SARS-CoV-2 receptor ACE2 by tobacco carcinogen-induced Skp2 in lung epithelial cells

**DOI:** 10.1101/2020.10.13.337774

**Authors:** Gui-Zhen Wang, Qun Zhao, Fan Liang, Chen Zhang, Hui Zhang, Jun Wang, Zhen-Yin Chen, Ran Wu, Hong Yu, Bei-Bei Sun, Hua Guo, Ruie Feng, Kai-Feng Xu, Guang-Biao Zhou

**Affiliations:** State Key Laboratory of Molecular Oncology, National Cancer Center/National Clinical Research Center for Cancer/Cancer Hospital, Chinese Academy of Medical Sciences and Peking Union Medical College, Beijing 100021, China; State Key Laboratory of Membrane Biology, Institute of Zoology, Chinese Academy of Sciences & University of Chinese Academy of Sciences, Beijing 100101, China; Hubei University of Medicine, Shiyan 442000, Hubei Province, China; Department of Pathology, Peking Union Medical College Hospital, Chinese Academy of Medical Sciences, Beijing 100730, China; Department of Pulmonary and Critical Care Medicine, Peking Union Medical College Hospital, Chinese Academy of Medical Sciences, Beijing 100730, China; Guizhou University School of Medicine, Guiyang 550025, Guizhou Province, China; School of Chinese Materia Medica, Beijing University of Chinese Medicine, No. 11, Bei San Huan Dong Lu, Beijing 100029, China

**Keywords:** SARS-CoV-2, tobacco smoke, benzo(a)pyrene, ACE2, Skp2

## Abstract

An unexpected observation among the COVID-19 pandemic is that smokers constituted only 1.4-18.5% of hospitalized adults, calling for an urgent investigation to determine the role of smoking in SARS-CoV-2 infection. Here, we show that cigarette smoke extract (CSE) and carcinogen benzo(a)pyrene (BaP) increase *ACE2* mRNA but trigger ACE2 protein catabolism. BaP induces an aryl hydrocarbon receptor (AhR)-dependent upregulation of the ubiquitin E3 ligase Skp2 for ACE2 ubiquitination. ACE2 in lung tissues of non-smokers is higher than in smokers, consistent with the findings that tobacco carcinogens downregulate ACE2 in mice. Tobacco carcinogens inhibit SARS-CoV-2 Spike protein pseudovirions infection of the cells. Given that tobacco smoke accounts for 8 million deaths including 2.1 million cancer deaths annually and Skp2 is an oncoprotein, tobacco use should not be recommended and cessation plan should be prepared for smokers in COVID-19 pandemic.

---

Severe acute respiratory syndrome coronavirus 2 (SARS-CoV-2), the pathogen of coronavirus disease 2019 (COVID-19), has infected 37,808,273 individuals and caused 1,080,790 deaths as of Oct 13, 2020 (Johns Hopkins University of Medicine Coronavirus Resource Center, 2020). The association between tobacco smoking and COVID-19 incidence and severity remains controversial (CDC COVID-19 Response Team, 2020; Farsalinos et al, 2020; Guan et al, 2020; Lippi & Henry, 2020; Miyara et al, 2020; Vardavas & Nikitara, 2020; World Health Organization, 2020a). Observational studies (CDC COVID-19 Response Team, 2020; Guan et al, 2020; Miyara et al, 2020) and meta-analyses (Farsalinos et al, 2020; Vardavas & Nikitara, 2020) showed that among the confirmed COVID-19 cases, smoker patients constituted a relatively small proportion (1.3% - 14.5%) of the total cases. Active smoking is not associated with severity of COVID-19 (Lippi & Henry, 2020), but is most likely associated with the negative progression and adverse outcomes of this disease (Vardavas & Nikitara, 2020). Based on these findings, pharmaceutical nicotine administration is suggested to be a therapeutic/preventive approach for SARS-CoV-2 infection (Miyara et al, 2020). Meta-analysis conducted by World Health Organization (WHO) showed that smokers constituted 1.4% - 18.5% of hospitalized adults, but smoking is associated with increased severity of the disease and death from COVID-19 (World Health Organization, 2020a). Therefore, the effects of tobacco smoking on SARS-CoV-2 infection should be determined by an experimental study.

## Methods

### Patient samples

The study was approved by the research ethics committees of the Chinese Academy of Medical Sciences Cancer Institute and Hospital. The lung biopsy samples of patients with benign diseases (Supplementary Table 1) were obtained from Department of Pathology, Peking Union Medical College Hospital, and were analyzed by immunohistochemistry assay using an anti-ACE2 and anti-Skp2 antibodies. Fresh normal lung tissues were collected 5 cm away from tumor samples of 49 previously untreated patients with lung adenocarcinoma (Supplementary Table 2), lysed, and tested by Western blot for the expression of ACE2 and Skp2.

### Animals

The animal studies were approved by the Institutional Review Board of the Chinese Academy of Medical Sciences Cancer Institute and Hospital. All animal studies were conducted according to protocols approved by the Animal Ethics Committee of our hospital. A/J mice (5-6 weeks old) were purchased from the Jackson Laboratory (Bar Harbor, Maine, USA), and were bred and maintained in a specific pathogen-free environment. The mice were exposed to cigarette smoke (Gebel et al, 2010) generated by DSI’s Buxco Smoke Generator (Buxco, NC, USA) inside a perspex box, at a frequency of 8 cigarettes per day, 5 days per week for 3 weeks, or were treated with BaP at 100 mg/kg or NNK at 50 mg/kg twice a week for 5 weeks (Wang et al, 2015b).

### Antibodies and reagents

Antibodies used in this study included rabbit polyclonal anti-human ACE2 (#ab15348, Abcam, Cambridge, MA, USA; 1:200 for immunofluorescence), rabbit monoclonal anti-human ACE2 (#ab108252, Abcam; 1:1000 for Western blot), rabbit polyclonal anti-human ACE2 (#4355, Cell Signaling Technology, Danvers, MA, USA; 1:1000 for Western blot), anti-TMPRSS2 (#ab92323, Abcam; 1:1000 for Western blot), rabbit anti-Furin (#ab183595, Abcam,; 1:1000 for Western blot), rabbit anti-Skp2 (#2652, Cell Signaling Technology; 1:1000 for Western blot), anti-GAPDH (#5174, Cell Signaling Technology; 1:1000 for Western blot), and anti-β-Actin (#A5441, Sigma, St. Louis, MO, USA; 1:5000 for Western blot). BaP (#B1760) and NNK (#78013) were purchased from Sigma. Cyclohexymide (CHX), epoxomicin, MG132, chloroquine (CQ), phorbol 12-myristate 13-acetate (PMA), azilsartan, candesartan, olmesartan, and eprosartan were purchased from Selleck (Shanghai, China). Cigarette smoke extract (CSE) was prepared as previously described (Carnevali et al, 2003).

### Cell culture

The cells were cultured in Dulbecco modified Eagle medium (DMEM) containing 10% fetal bovine serum (FBS; Gibco/BRL, Grand Island, NY), 100 U/ml penicillin, 100 mg/ml streptomycin, and were treated with CSE, BaP, NNK, CHX, MG132, epoxomicin, and CQ as indicated (Liu et al, 2014; Wang et al, 2019; Wang et al, 2015a). The cells were transfected with 50 nM double-stranded siRNA oligonucleotides (Supplementary Table 3) using HiPerFect Transfection Reagent (Qiagen, Crawley, UK).

The total RNA was isolated using the TRIZOL Reagent (Invitrogen, Frederick, MD, USA) according to the manufacturer’s instruction. Total RNA (2 μg) was annealed with random primers at 65 °C for 5 min. The cDNA was synthesized using a 1st-STRAND cDNA Synthesis Kit (Fermentas, Pittsburgh PA, USA). Quantitative RT-PCR was carried out using SYBR Premix ExTaq^™^ (Takara Biotechnology, Dalian, China). The primers used in this study were listed in Supplementary Table 3.

### Western blotting

Lung tissue samples and the cells were lysed on ice for 30 min in RIPA buffer (50 mM Tris-HCl pH 7.4, 150 mM NaCl, 0.1% SDS, 1% deoxycholate, 1% Triton X-100, 1 mM EDTA, 5 mM NaF, 1 mM sodium vanadate, and protease inhibitors cocktail), quantitated, and subjected to 10-15% SDS-PAGE, electrophoresed and transferred to a nitrocellulose membrane. After blocking with 5% non-fat milk in Tris-buffered saline, the membrane was washed and incubated with indicated primary and secondary antibodies and detected by Luminescent Image Analyzer LSA 4000 (GE, Fairfield, CO, USA). Densitometry analyses of immunoblot bands were performed to quantitate the expression level of the proteins.

### Immunohistochemistry analysis

The formalin-fixed, paraffin-embedded human or mouse lung cancer tissue specimens (5 μm) were deparaffinized through xylene and graded alcohol, and subjected to a heat-induced epitope retrieval step in citrate buffer solution. The sections were then blocked with 5% BSA for 30 min and incubated with indicated antibodies at 4 °C overnight, followed by incubation with secondary antibodies for 90 min at 37 °C. Detection was performed with 3, 3’-diaminobenzidine (DAB, Zhongshan Golden Bridge Biotechnology Co., Ltd, Beijing, China) and counterstained with hematoxylin, dehydrated, cleared and mounted using the routine processing. The immunoreactivity score was calculated as IRS (0–12)=RP (0–4) × SI (0–3), where RP is the percentage of staining-positive cells and SI is staining intensity.

### *In vitro* ubiqutination assay

Ubiquitination assay was performed with Ubiquitylation Assay Kit (abcam; ab139467) using recombinant carrier free ACE2 (# 933-ZN-010) and Skp1/Skp2 (#E3-521-025) proteins bought from R&D Systems (Minneapolis, MN, USA) or Flag-ACE2 and Flag-Skp1/Skp2 purified from 293T cells. The proteins were incubated with 50 μl reaction mixture (Boston Biochem), 2 mM Mg-ATP (abcam), 0.5 μg E1 (abcam), 2.5 μg UbcH5a/UBE2D1 (Boston Biochem), and 2.5 μg ubiquitin (abcam) at 37°C for 2 h. After incubation, the reaction was terminated with SDS loading buffer, and the ubiquitination was detected using Western blot assays.

### SARS-CoV-2 Spike protein pseudovirions

Pseudovirions were purchased from Sino Biological Inc. (Beijing, China), and were produced by transfection of 293T cells with psPAX2, pLenti-GFP, and plasmids encoding SARS-CoV-2 Spike protein using polyetherimide. To infect 16HBE and *293T-ACE2* cells with pseudovirions, the cells were seeded onto 96-well plates, pre-treated with CSE (10%), BaP (5 μM), or NNK (5 μM) for 48 h, followed by co-incubation with 100 μl media containing pseudovirions for 48 h. The cells were lysed with 60 μl medium containing 50% Steady-glo (promega) and measured by quantification of the luciferase activity using a Multi-Mode Reader (BioTek, Sunnyvale, CA, USA).

### Statistical analysis

All statistical analyses were conducted using the software SPSS 26.0 for Windows (Chicago, IL, USA). Statistically significant differences were determined by Students *t*-test or Fisher’s exact test, and correlation between relative ACE2 and Skp2 expression levels was tested by Spearman correlation analysis. *P* values less than 0.05 were considered statistically significant in all cases.

## Results

### Tobacco smoke induces dual effects on ACE2

SARS-CoV-2 enters target cells by binding to its receptor ACE2 and using the serine protease TMPRSS2 for viral spike (S) protein priming (Hoffmann et al, 2020; Wang et al, 2019; Zou et al, 2020). We tested the effects of cigarette smoking on *ACE2* expression, and reported that treatment of human normal lung epithelial 16HBE cells with cigarette smoke extract (CSE) (Wang et al, 2019) at final concentrations of 5% to 10% slightly upregulated *ACE2* at mRNA level (Figure 1A), consistent with previous analyses of transcriptomic datasets (Cai et al, 2020; Smith et al, 2020). We then tested the effects of tobacco smoking on ACE2 at protein level. Unexpectedly, we found that treatment of the cells with CSE at 5% to 20% for 48 h (Figure 1B) or at 10% for 48 to 72 h (Figure 1C) significantly downregulated ACE2 at protein level, revealed by Western blot assay and densitometry analysis of immunoblot bands. The discrepancy in changes in mRNA and protein levels demonstrated the complicated effects of tobacco smoking on this receptor, and modulation of ACE2 protein may underlie the effect of smoking on SARS-CoV-2 infection.

**Figure 1.**
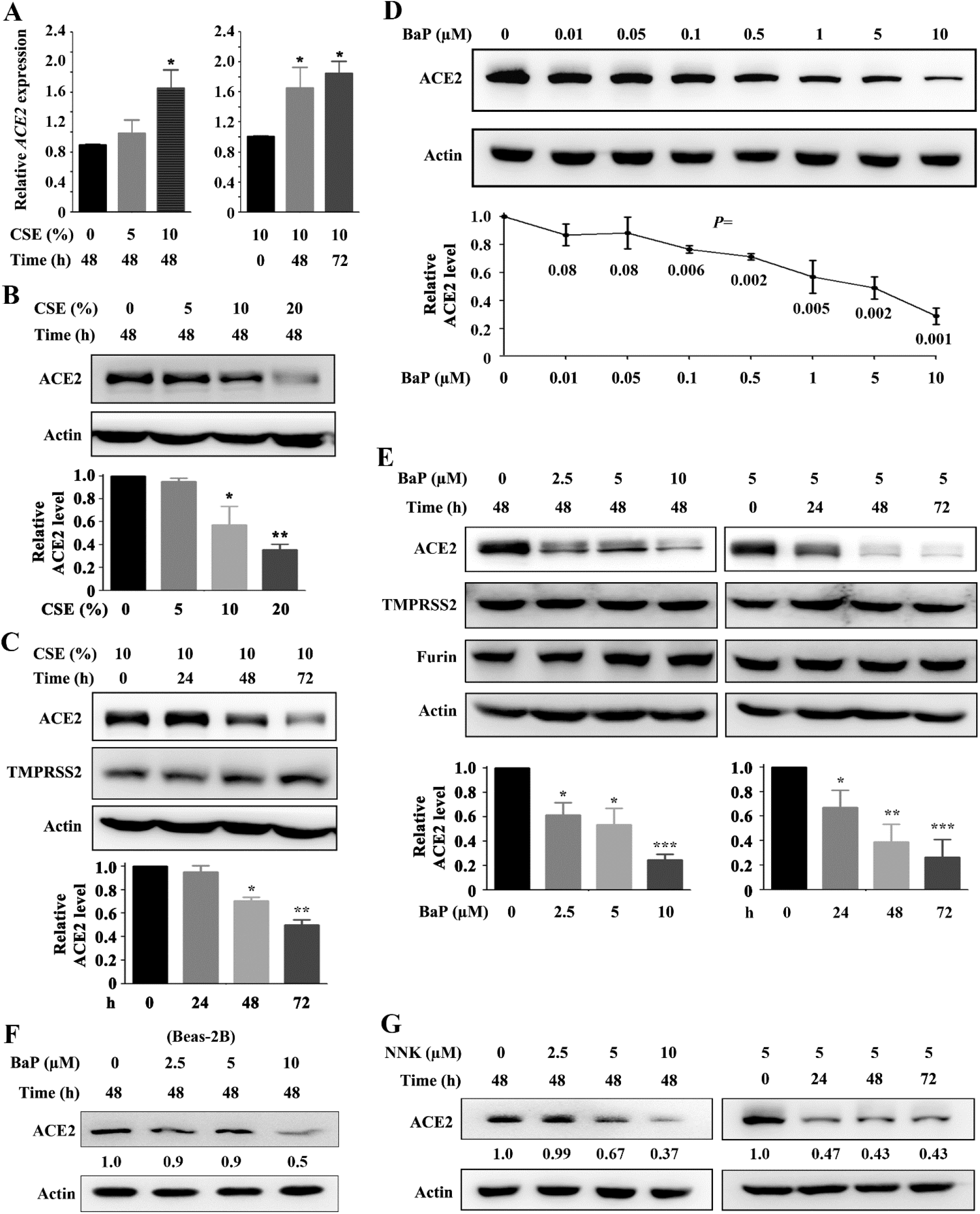
Effects of tobacco carcinogens on ACE2 expression in lung epithelial cells. (A) 16HBE cells were treated with CSE at indicated protocols, lysed, RNA was extracted, and *ACE2* expression at mRNA level was evaluated by quantitative reverse transcription-polymerase chain reaction (RT-PCR). *P* values, Student’s *t*-test. *, *P* < 0.05. (B, C) 16HBE cells were treated with CSE at indicated protocols, and ACE2 expression at protein level was detected by Western blot. The relative ACE2 levels were determined by densitometry analysis of immunoblot bands and normalized to Actin expression levels. *P* values, Student’s *t*-test. \**P* < 0.05, \*\**P* < 0.01. (D, E) 16HBE cells were treated with BaP at indicated concentrations for 48 h (D) or at indicated protocols (E), lysed, and subjected to Western blot using indicated antibodies. The relative ACE2 levels were determined as described above. *P* values, Student’s *t*-test. \**P* < 0.05, \*\**P* < 0.01, ***, *P* < 0.001. (F) Beas-2B cells were treated with BaP, and ACE2 expression at protein level was detected by Western blot using cell lysates and indicated antibodies. Numbers under the Western blot bands are the relative expression values to Actin determined by densitometry analysis. (G) 16HBE cells were treated with NNK, lysed, and subjected to Western blot using indicated antibodies. Numbers under the Western blot bands are the relative ACE2 expression values to Actin.

### Tobacco carcinogens trigger ACE2 degradation

The effects of several representative components of tobacco smoking on ACE2 were tested, and the results showed that treatment of 16HBE cells with nicotine at 2.5 to 10 μM for 48 h or at 5 μM for 24 to 72 h did not perturb ACE2 expression (Supplementary Figure 1A). Of note, group 1 carcinogen (carcinogenic to humans) benzo(a)pyrene (BaP) induced downregulation of ACE2 at 0.01 to 10 μM treatment for 48 h (Figure 1D). Treatment of 16HBE cells with BaP at 2.5 to 10 μM for 48 h or at 5 μM for 24 to 72 h significantly downregulated ACE2 expression (Figure 1E). CSE and BaP did not affect the expression of TMPRSS2 or Furin (Figure 1C, E). BaP downregulated ACE2 in normal human lung epithelial Beas-2B cells (Figure 1F), human alveolar epithelial A549 cells, and human embryonic kidney 293T cells transfected with pCMV-*ACE2*-hygro plasmid (Supplementary Figure 1B). In macrophages derived from human monocytic cell line THP-1 by stimulation with phorbol 12-myristate 13-acetate (PMA), the expression of ACE2 was low and BaP did not perturb its expression (Supplementary Figure 1C) but upregulated cytokines *interferon-γ* (*IFNγ*), *interleukin-1α* (*IL-1α*), *IL-1β, IL-2, IL-6*, and *tumor necrosis factor α* (*TNFα*) that are associated with cytokine storm in COVID-19 (Zhou et al, 2020). Moreover, another tobacco group 1 carcinogen nitrosamine 4-(methylnitrosamino)-1-(3-pyridyl)-1-butanone (NNK) induced downregulation of ACE2 in 16HBE cells (Figure 1G). On the contrary, the angiotensin receptor blockers azilsartan, candesartan, olmesartan but not eprosartan, induced upregulation of ACE2 (Supplementary Figure 2A), consistent with previous reports (Gheblawi et al, 2020). These compounds did not perturb the expression of TMPRSS2 or Furin in the cells (Supplementary Figure 2B).

### Tobacco carcinogen induces ubiquitination and degradation of ACE2

BaP was used as a representative compound to dissect the mechanism of tobacco smoking-induced ACE2 downregulation at protein level. We showed that BaP treatment at 2.5 – 5 μM for 48 h or at 5 μM for 48 – 72 h slightly increased *ACE2* mRNA level (Figure 2A). BaP increased *ACE2* promoter-driven luciferase activity (Figure 2B). We tested ACE2 protein stability by incubation of 16HBE cells with protein synthesis inhibitor cycloheximide (CHX), and found that treatment of the cells with CHX (50 μg/mL) for 12 h did not cause significant downregulation of ACE2 (Figure 2C), suggesting that this receptor was quite stable. However, combinatorial use of CHX and BaP led to downregulation of ACE2 at 6 to 12 h, revealed by Western blot and densitometry analysis of protein bands (Figure 2C). These results suggested that BaP may induce proteolytic degradation of ACE2. Proteasome and lysosome are two critical sites for degradation of ubiquitinated substrate proteins, and we found that proteasome inhibitors epoxomicin and MG132 (Figure 2D) and lysosome inhibitor chloroquine (CQ; Figure 2E) inhibited BaP-induced ACE2 downregulation. In cells transfected with hemagglutinin-ubiquitin (HA-Ub) plasmid (Kamitani et al, 1997), BaP induced downregulation of ACE2 and enhanced Ub-ACE2 binding affinity (Figure 2F). In 16HBE cells treated with 5 μM for 24 h, BaP downregulated ACE2 and potentiated ubiquitination of ACE2 (Figure 2G).

**Figure 2.**
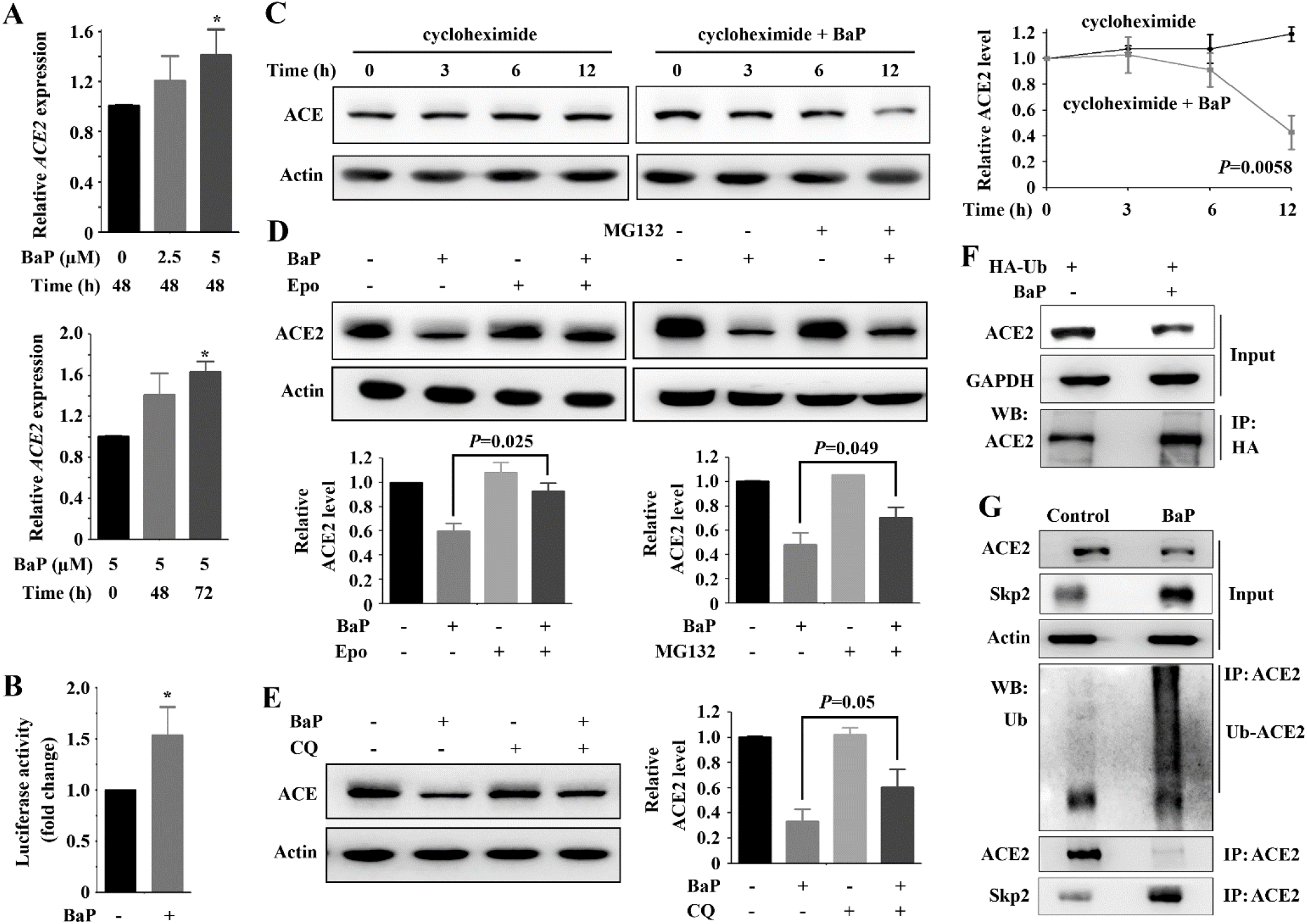
BaP induces ACE2 degradation. (A) 16HBE cells were treated with BaP, lysed, RNA was extracted, and *ACE2* expression at mRNA level was evaluated by quantitative RT-PCR. *P* values, Student’s *t*-test. *, *P* < 0.05. (B) The HLF cells were transfected with *ACE2* promoter-luciferase reporter construct, treated with BaP for 48 h, and assessed by the luciferase assay. (C) 16HBE cells were treated with cycloheximide (50 μg/mL) in the absence or presence of BaP (5 μM), lysed, and subjected to Western blot assays. The relative ACE2 levels were determined as described above. *P* values, Student’s *t*-test. (D) 16HBE cells were pre-treated with BaP (5 μM) for 24 h, subsequently co-incubated with epoxomicin (10 μM) or MG132 (10 μM) for 12 h, and then lysed for Western blot analyses. (E) 16HBE cells were treated with chloroquine (CQ; 40 μM) and BaP (5 μM) for 36 h, and lysed for Western blot analyses. (F) 293T cells were transfected with HA-Ub for 24 h, treated with BaP (5 μM) for additional 48 h, lysed, and subjected to immunoprecipitation and immunoblot using indicated antibodies. Ub, ubiquitin. (G) 16HBE cells were treated with BaP (5 μM) for 48 h, lysed, and subjected to immunoprecipitation and immunoblot using indicated antibodies.

### Skp2 mediates carcinogen-induced ACE2 ubiquitination and degradation

To identify the E3 ubiquitin ligase that mediates BaP-induced ACE2 degradation, the expression of several critical E3 ligases, e.g., the F-box (40 amino acid motif)-containing E3 ligase S-phase kinase associated protein 2 (Skp2), β-transducin repeat containing E3 ubiquitin protein ligase (β-TRCP), the nuclear interaction partner of ALK (NIPA), the RING finger and WD repeat domain 3 (RFWD3), and deubiquitinase ubiquitin-specific protease 10 (USP10) were tested in cells treated with BaP. We showed that upon BaP treatment, the expression of Skp2 (Zhang et al, 1995) was drastically upregulated at protein (Figure 3A) and mRNA (Figure 3B) levels. However, the expression of β-TRCP, NIPA, RFWD3, USP10, and Skp1 which is a core subunit of the Skp1-Cul1-F-box protein (SCF) complex, was not significantly perturbed by BaP treatment (Figure 3A).

**Fig. 3.**
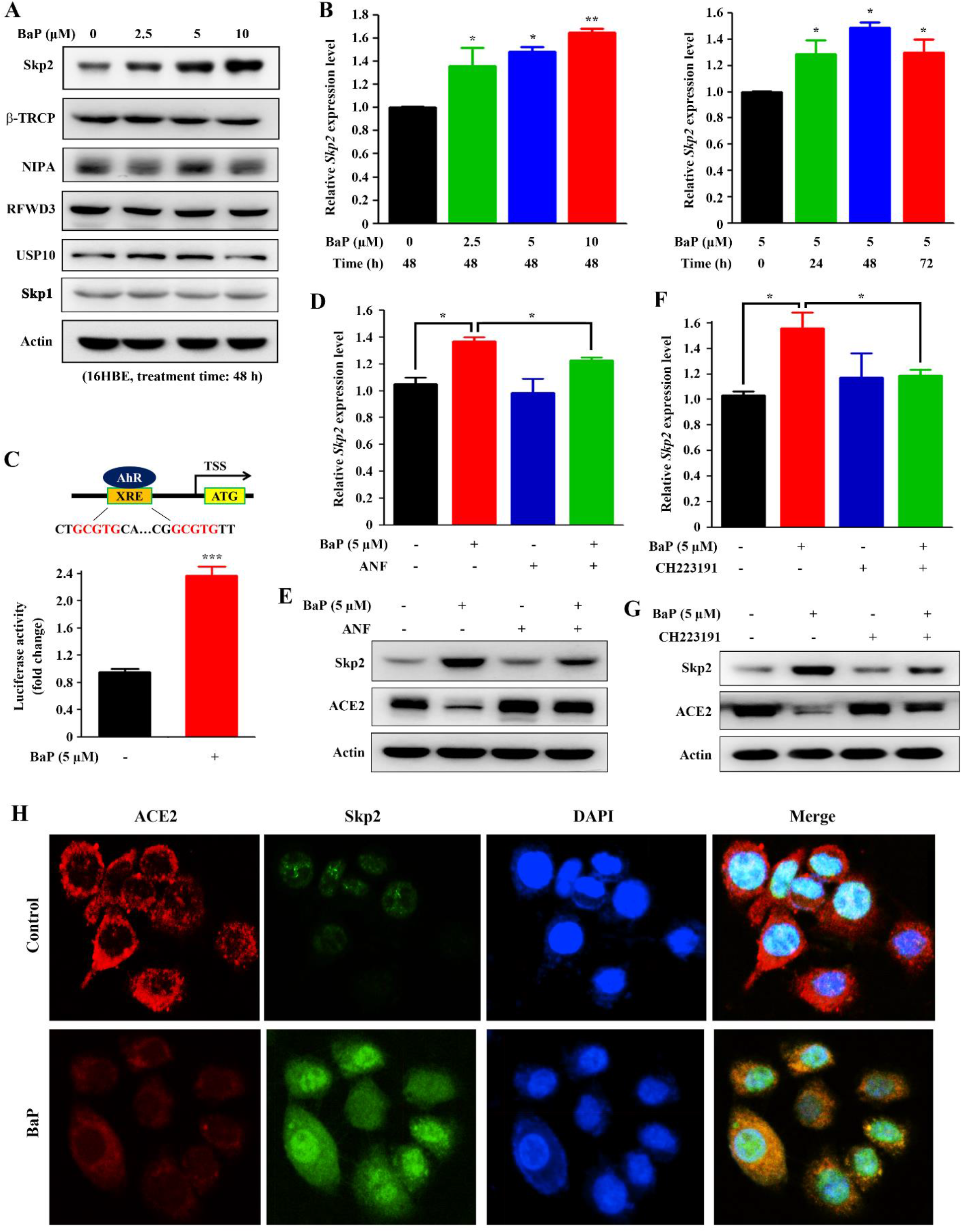
BaP induces an AhR-dependent Skp2 upregulation. (A) 16HBE cells were treated with BaP at indicated concentrations for 48 h, lysed, and subjected to immunoblot using indicated antibodies. (B) 16HBE cells were treated with BaP as indicated, lysed, RNA were extracted, and analyzed by quantitated RT-PCR. Student’s *t* test, *, P < 0.05; **, P < 0.01. Error bars, SD. (C) *ACE2* is a target gene of AhR. The upper panel shows the AhR binding site of *Skp2* promoter. TSS, transcription start site. Lower panel, 16HBE cells were transfected with the XRE element of *Skp2* promoter-luciferase reporter construct, treated with BaP for 48 h, and assessed by the luciferase assays. Student’s *t* test, ***, P < 0.001. Error bars, SD. (D - F) 16HBE cells were treated with BaP in the absence or presence of AhR antagonist ANF (D, E) or CH223191 (F, G), lysed, and analyzed by quantitative RT-PCR (D, F) or Western blot (E, G). Student’s *t* test, *, P < 0.05. Error bars, SD. (H) The expression of ACE2 and Skp2 in 16HBE cells treated with vehicle control (DMSO) or BaP at 10 μM for 48 h. The cells were harvested and detected by immunofluorescence assays using indicated antibodies.

The helix-loop-helix transcription factor aryl hydrocarbon receptor (AhR) plays a key role in the regulation of biological responses to polycyclic aromatic hydrocarbons (PAHs) and dioxin (Tsay et al, 2013). By analyzing its promoter sequence, we found that *Skp2* harbors two xenobiotic-responsive elements (XRE), 5’-<u>GCGTG</u>-3’ and 5’-<u>GCGTG</u>-3’, in its promoter region (Figure 3C), indicating that it is a target gene of AhR. We showed that BaP increased *Skp2* promoter-driven luciferase activity (Figure 3C), and AhR inhibitors alpha-naphthoflavone (ANF; Figure 3D, E) and CH223191 (Figure 3F, G) suppressed BaP-induced Skp2 upregulation at both mRNA (Figure 3D, F) and protein (Figure 3E, G) levels, and inhibited BaP-induced ACE2 downregulation (Figure 3D-G).

We showed that in 16HBE cells upon BaP treatment, Skp2 upregulation was concomitantly detected with downregulation of ACE2, and co-localization of ACE2 and Skp2 was seen within the cells (Figure 3H). In consistent with these observations, ACE2 expression was negatively associated with Skp2 expression in several cell lines (Supplementary Figure 3). By co-immunoprecipitation assay, we reported that Skp2 could bind to ACE2 but not RFWD3 in 16HBE and pCMV-*ACE2*-transfected 293T (Figure 2G, Figure 4A) cells in the absence and presence of BaP treatment, and BaP treatment enhanced ACE2-Skp2 binding affinity. By expression and purification of truncated Skp2 mutants (Figure 4B), we showed that two domains (183 – 292 and 293 – 424 amino acids) in C-terminal of Skp2 were responsible for interaction with ACE2, whereas the N-terminal (1 – 182 amino acids) of this E3 ligase was unable to bind ACE2 (Figure 4C). Exogenous Skp2 also bound ACE2 and caused enhanced ubiquitination and downregulation of ACE2 in 16HBE and 293T-ACE2 cells (Figure 4D). By CHX chase experiment, we showed that ectopic expression of Skp2 induced decrease of ACE2 stability in 16HBE cells (Figure 4E). Meanwhile, knockdown of Skp2 by specific small interfering RNA (siRNA) suppressed BaP-induced proteolytic degradation as well as ubiquitination of ACE2 (Figure 4F, G). Silencing of Skp2 by si*Skp2* attenuated BaP-induced ACE2 ubiquitination and degradation in 293T-*ACE2* cells (Figure 4H). To further determine whether Skp2 could induce ACE2 ubiquitination, an *in vitro* ubiquitination assay was performed using Flag-ACE2 and Flag-Skp2 proteins purified from 293T cells, and the results showed that Skp2 was able to trigger ACE2 ubiquitination (Figure 4I). In *in vitro* ubiquitination assay using recombinant carrier free ACE2 and Skp2 proteins bought from R&D Systems, we confirmed that Skp2 was able to trigger ACE2 ubiquitination (Figure 4J), demonstrating that Skp2 is the E3 ligase for ACE2 ubiquitination and degradation.

**Figure 4.**
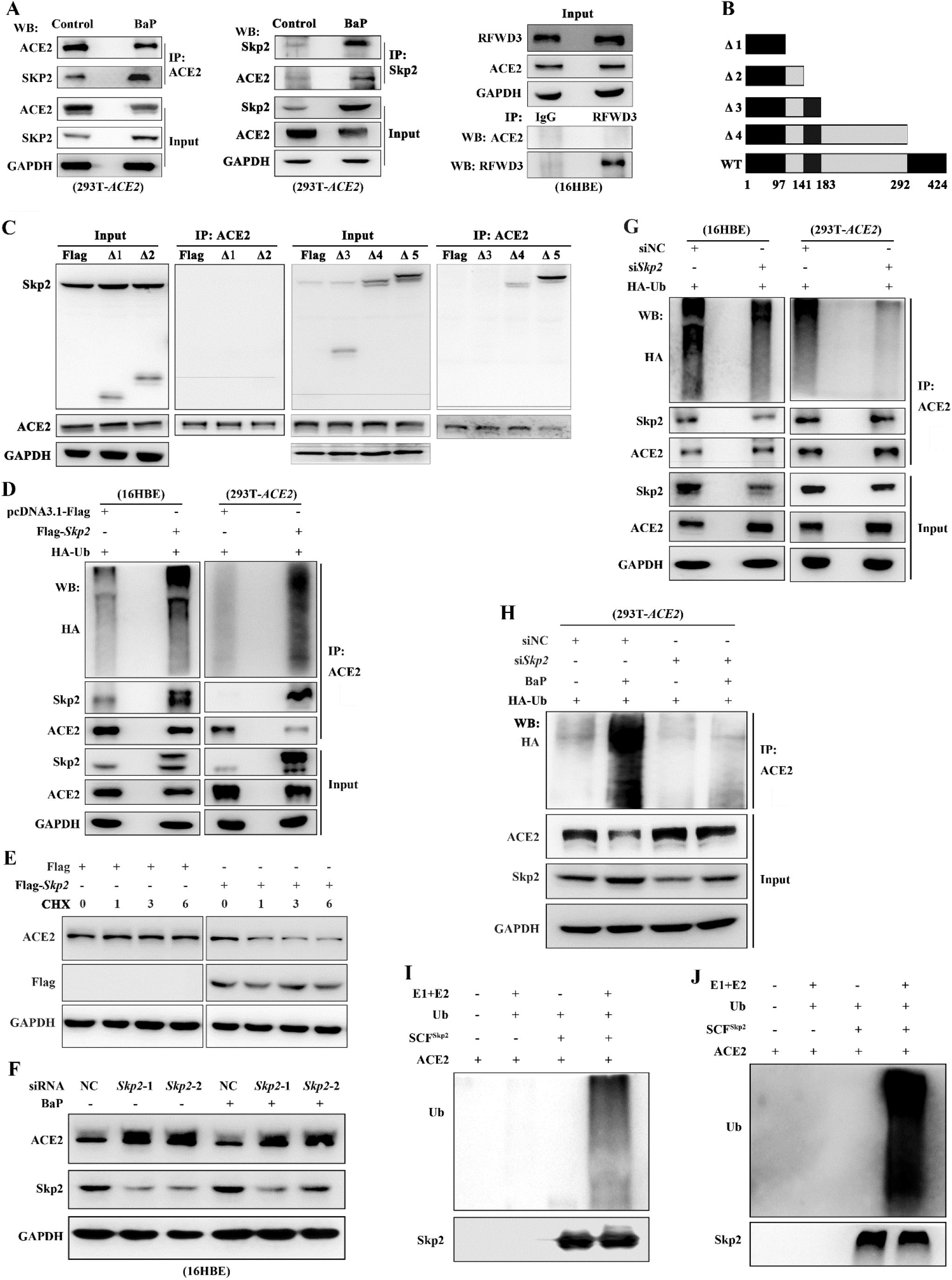
Skp2 mediates ubiquitination and degradation of ACE2 induced by BaP. (A) *293T-ACE2* cells were treated with BaP (5 μM) for 48 h, lysed, and subjected to immunoprecipitation and immunoblot using indicated antibodies. Lysates of 16HBE cells were subjected to immunoprecipitation and immunoblot assays using indicated antibodies (right panel). (B, C) Schematic representation of *Skp2* truncated mutants (B), which were transfected into 293T-*ACE2* cells for protein purification and subsequent immunoblotting assays using indicated antibodies (C). (D) 16HBE and 293T-*ACE2* cells were transfected with Flag-*Skp2* and HA-Ub for 48 h, lysed, and subjected to immunoprecipitation and immunoblot using indicated antibodies. (E) 16HBE cells were transfected with Flag-*Skp2*, treated with CHX, lysed, and subjected to immunoblot using indicated antibodies. (F) 16HBE cells were transfected with si*Skp2* for 24 h, followed by co-incubation with BaP (5 μM) for 48 h, lysed, and subjected to immunoblot using indicated antibodies. (G) 16HBE and 293T-*ACE2* cells were transfected with si*Skp2* and HA-Ub for 48 h, lysed, and subjected to immunoprecipitation and immunoblot using indicated antibodies. (H) 293T-*ACE2* cells were transfected with si*Skp2* and HA-Ub for 48 h, followed by treatment with BaP at 5 μM for additional 48 h. The cells were lysed, and subjected to immunoprecipitation and immunoblot using indicated antibodies. (I) *In vitro* ubiquitination assay using Flag-SCF^Skp2^ and Flag-ACE2 purified from 293T cells. (J) *In vitro* ubiquitination assay using recombinant carrier free ACE2 and SCF^Skp2^ proteins bought from R&D Systems.

### Tobacco carcinogens induced ACE2 downregulation *in vivo*

To test the *in vivo* effect of tobacco smoking on ACE2 expression, immunohistochemistry (IHC) assay was performed on lung biopsy samples of patients with benign diseases (Supplementary Table 1). The results showed that ACE2 was expressed on membrane and cytoplasm of the cells, and smokers had much lower ACE2 than non-smokers (Figure 5A, B). On the contrary, the expression of Skp2 was low in non-smokers and high in smokers (Figure 5A, B), showing a negative correlation trend with ACE2 in the patients. In another setting, Western blot analysis was performed using lysates of specimens harvested from normal lung tissues 5 cm away from tumor tissues of patients with lung adenocarcinoma (Supplementary Table 2), and we reported that in smoker patients the expression of ACE2 was lower whereas Skp2 was higher than those in non-smoker patients (Figure 5C, D). The relationship between ACE2 and Skp2 expression levels was analyzed by Spearman analysis, and a negative correlation between the two variables was detected (Figure 5E).

**Figure 5.**
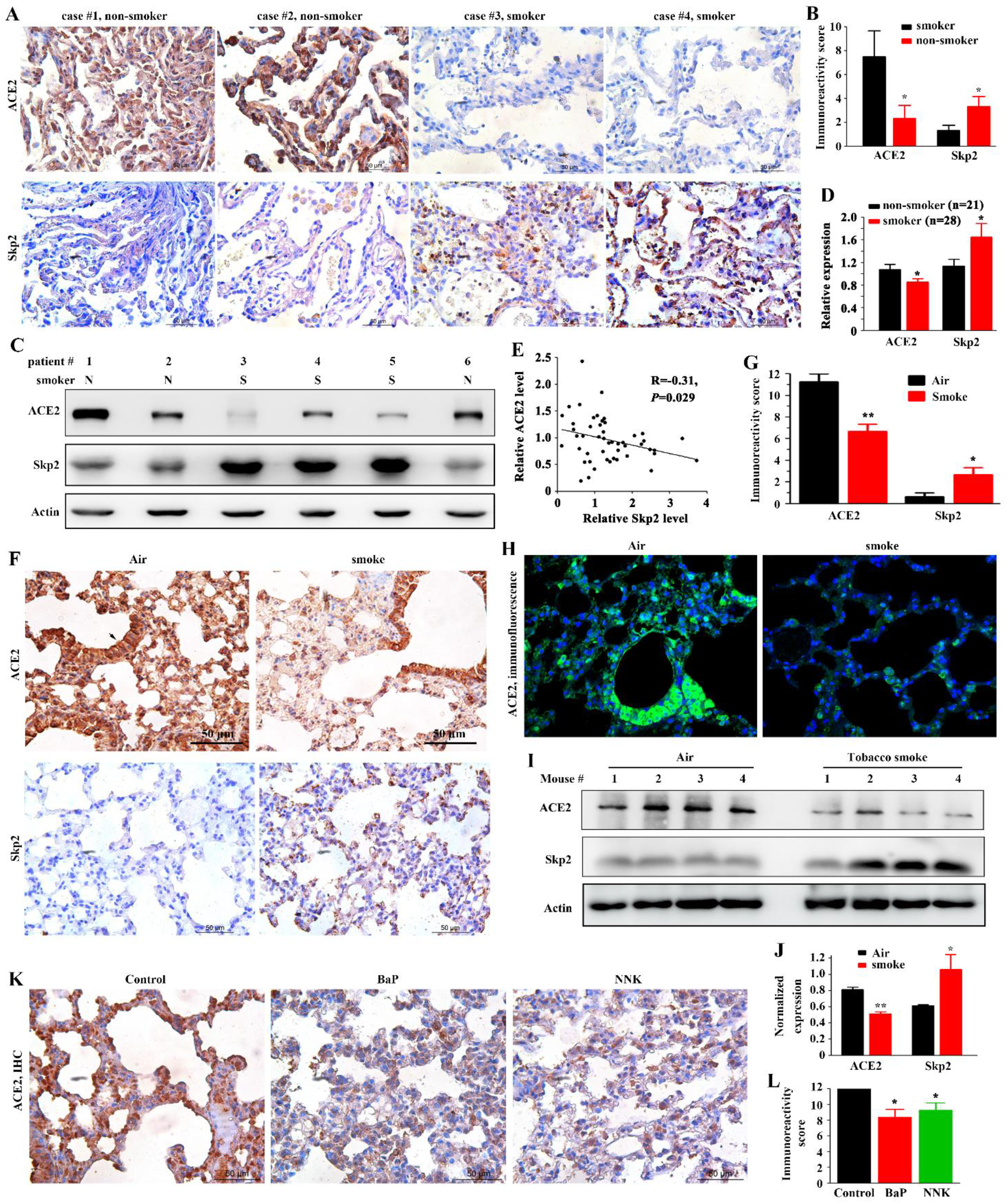
Tobacco smoke and BaP induce downregulation of ACE2 *in vivo*. (A, B) Immunohistochemistry analysis of lung biopsy samples of patients with benign disease (A), and immunoreactivity score of ACE2 and Skp2 was calculated (B). (C, D) Western blot assays of normal lung tissues harvested 5 cm away from tumor tissues of patients with lung adenocarcinoma (C) and quantification of ACE2 and Skp2 determined by densitometry analyses of immunoblot bands (D). (E) Spearman correlation analysis of ACE2 and Skp2 relative expression levels of the patients using the results of (D). (F, G) Immunohistochemistry analyses of lung tissues of mice exposed to clean air or tobacco smoke (F), and immunoreactivity score of ACE2 and Skp2 was calculated (G). Arrow, ciliated cells also express ACE2. (H) Immunofluorescence assays of lung tissues of mice exposed to clean air or tobacco smoke. (I, J) Western blot assays of lung tissues of mice exposed to clean air or tobacco smoke (I) and quantification of ACE2 and Skp2 determined by densitometry analyses of immunoblot bands (J). (K, L) Immunohistochemistry analyses of lung tissues of mice treated with vehicle control, BaP, or NNK (K). Immunoreactivity score of ACE2 was calculated (L).

To confirmed these observations, the A/J mice were exposed to cigarette smoke with filtered conditioned air of 750 μg total particulate matter per liter for 20 days (Wang et al, 2019), and lung tissues of the mice were stained with anti-ACE2 and anti-Skp2 antibodies. IHC assays (Figure 5F) and related IRS (Figure 5G) showed that ACE2 was lower while Skp2 was higher in mice exposed to tobacco smoking than in mice exposed to clean air. These results were confirmed by immunofluorescence (Figure 5H) and Western blot/related densitometry assays (Figure 5I, J) of these lung tissues. In mice treated with BaP at 100 mg/kg twice a week for 5 weeks, the ACE2 expression in lung tissues was decreased compared with that in mice treated with vehicle control (Figure 5K, L). Treatment with NNK at 50 mg/kg twice a week for 5 weeks also caused downregulation of ACE2 in lung tissues of the mice (Figure 5K, L).

### Effects of tobacco carcinogens on SARS-CoV-2 Spike protein pseudovirus infection

The effects of CSE on viral entry were tested in 16HBE cells using the SARS-CoV-2 spike (S) protein pseudovirus (Ou et al, 2020), and the results showed that while the pseudovirion was able to enter into the cells, treatment with CSE at 10% for 48 h reduced the entry of pseudovirus by approximately 25%, revealed by relative luciferase activity (Figure 6A). BaP (5 μM) and NNK (5 μM) inhibited SARS-CoV-2 S protein pseudovirus from entry into the cells by 35% and 30%, respectively (Figure 6A). In 293T cells transfected with pCMV-ACE2-hygro plasmid, treatment with CSE, NNK, and BaP reduced pseudovirus entry by 15%, 30%, and 40%, respectively (Figure 6B).

**Figure 6.**
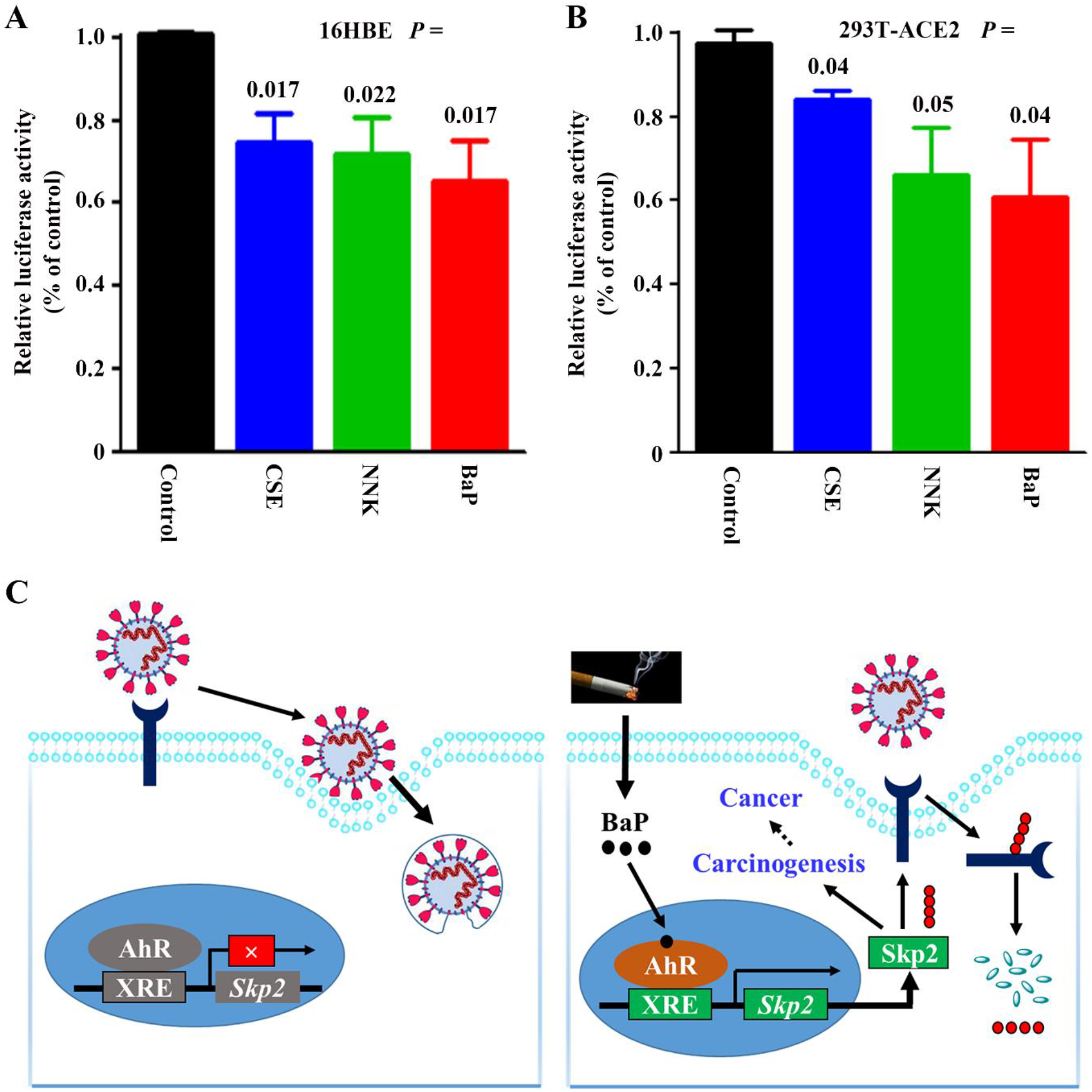
Inhibition of SARS-CoV-2 S protein pseudoviron entry by CSE, BaP, and NNK. (A) Entry of SARS-CoV-2 S protein pseudovirions into 16HBE cells treated with indicated agents. (B) Entry of SARS-CoV-2 S protein pseudovirions into 293T-*ACE2* cells treated with indicated agents. (C) Schematic representation of tobacco smoke-induced ACE2 degradation.

## Discussion

Tobacco smoke accounts for more than 8 million deaths each year worldwide, with 7 million deaths from direct tobacco use and 1.2 million deaths of non-smokers being exposed to second-hand smoke (World Health Organization, 2020b). There is strong evidence for a causal association between smoking and 29 diseases (14 types of cancers, 7 categories of cardiovascular disease, 3 categories of respiratory disease, diabetes, renal failure, infections, intestinal ischemia, and liver cirrhosis) (Carter et al, 2015). Tobacco and smoking contain more than 70 carcinogens and cause cancers via complicated mechanisms (Centers for Disease Control and Prevention (US) et al, 2010; Zhou, 2019). Therefore, great efforts should be made to reduce tobacco use and help smokers to quit.

Effective drugs for COVID-19 remain rare, and vaccines for SARS-CoV-2 are still under investigation (Zhou et al, 2020). CSE inhibits SARS-CoV-2 S protein pseudovirus infection, supporting the clinical findings that smokers might have a lower probability of developing SARS-CoV-2 infection as compared to the general population (Miyara et al, 2020; Vardavas & Nikitara, 2020). Among the tobacco compounds, BaP and NNK, but not nicotine, inhibit SARS-CoV-2 S protein pseudovirion from infection of the cells. These results do not suggest an efficacy of nicotine in prevention/treatment of COVID-19. Both BaP and NNK are notorious carcinogens that cause comprehensive hazardous effects on humans and induce carcinogenesis (Zhou, 2019). BaP also induces upregulation of Skp2, an oncoprotein that plays a pivotal role in cell cycle progression and proliferation (Liu et al, 2020). Therefore, the generalized advice to quit smoking as a measure to improve health risk remains valid, because it is the carcinogens that are responsible for ACE2 degradation.

ACE2 is a glycoprotein metalloprotease that exists in membrane-bound and soluble forms and has enzymatic and non-enzymatic functions. It is expressed in type II alveolar cells, myocardial cells, proximal tubule cells of kidney, ileum and esophagus epithelial cells, bladder urothelial cells, and other types of cells (Zou et al, 2020). We showed that tobacco carcinogens exerts dual effects on ACE2, i.e., upregulation at mRNA and downregulation at protein levels, in normal human lung epithelial cells. This discrepancy was consistent with studies into mRNA-protein relationship showing poor correlation between the two kinds of molecules (Xu et al, 2020). A study reports that angiotensin II (Ang-II) treatment induces ACE2 internalization into lysosomes where it is degraded (Deshotels et al, 2014). We show that lysosome is partially responsible for BaP-triggered ACE2 degradation, because lysosome inhibitor chloroquine partially rescues ACE2 from BaP-induced degradation. Proteasome also has a role in ACE2 catabolism, since epoxomicin and MG132 partially suppress ACE2 degradation. The fact that inhibition of either proteasome or lysosome can partially but not completely blocked the degradation of ACE2 suggests that the both organelles are critical to ACE2 catabolism, and the role of other forms of post-translational modification (e.g., phosphorylation and glycosylation) in ACE2 catabolism warrants further investigation. Currently, how NNK downregulates ACE2 remains unclear. We show that BaP induces a significant up-regulation of Skp2, which interacts with ACE2 and induces ubiquitination and degradation of the substrate. Therefore, our results partially unveil the mechanisms of action of tobacco carcinogens on ACE2 (Figure 6C). This receptor is required to maintain lung and cardiovascular functions (Gheblawi et al, 2020), whether ACE2 proteolysis is responsible for severity of patients with COVID-19 needs to be determined in the future studies.

## Supplementary information

Supplementary information contains 3 supplementary tables and 3 supplementary figures and legends.

## Acknowledgements

This work was jointly supported by the National Natural Science Funds for Distinguished Young Scholar (81425025), the Key Project of the National Natural Science Foundation of China (81830093), the CAMS Innovation Fund for Medical Sciences (CIFMS; 2019-I2M-1-003), and the National Natural Science Foundation of China (81672765, 81802796).

## Author contributions

The project was conceived by G.B.Z.. The experiments were designed by G.B.Z.. The experiments were conducted by G.Z.W., Q.Z., F.L., C.Z., Z.Y.C., R.W., H.Y., B.B.S. and G.H.. Biospecimens/materials were harvested/provided by H.Z., J.W., R.F., and K.F.X.. Data were analyzed by G.B.Z. and G.Z.W.. The manuscript was written by G.B.Z..

## Declarations of interests

No potential conflicts of interest were disclosed.

## Supplementary materials

**Supplementary Table 1.** Baseline demographic characteristics of the patients whose samples were analyzed by immunohistochemistry assay.

| Case # | Age<br>(year) | Sex | Smoking<br>status | Histological diagnosis |
| --- | --- | --- | --- | --- |
| 1 | 70 | Male | Never | Pulmonary chronic inflammation |
| 2 | 73 | Male | Never | Pulmonary chronic inflammation |
| 3 | 56 | Male | Never | Pulmonary chronic inflammation |
| 4 | 53 | Male | Never | Pulmonary chronic inflammation |
| 5 | 36 | Male | Never | Proliferation of alveolar epithelia and lymphoid tissue with carbon sedimentation |
| 6 | 63 | Male | Never | Fibrotic tubercle with carbon sedimentation |
| 7 | 47 | Male | Smoker | Pulmonary chronic inflammation |
| 8 | 33 | Male | Smoker | Pulmonary chronic inflammation |
| 9 | 51 | Male | Smoker | Pulmonary chronic inflammation |
| 10 | 50 | Male | Smoker | Inflammatory cell infiltration and cellulose exudation |
| 11 | 56 | Male | Smoker | Pulmonary carbon sedimentation and aggregation of phagocyte |
| 12 | 60 | Male | Smoker | Pulmonary chronic inflammation |

**Supplementary Table 2.** Baseline demographic characteristics of the patients with lung adenocarcinoma whose samples were detected by Western blot analyses.

| Characteristics | Total, n=49 | Smoker, n=28 | Non-smoker, n=21 |
| --- | --- | --- | --- |
| Gender |  |  |  |
| Male | 35 | 28 | 7 |
| Female | 14 | 0 | 14 |
| Age, year |  |  |  |
| < 65 | 38 | 21 | 17 |
| ≥ 65 | 11 | 7 | 4 |

**Supplementary Table 3.**
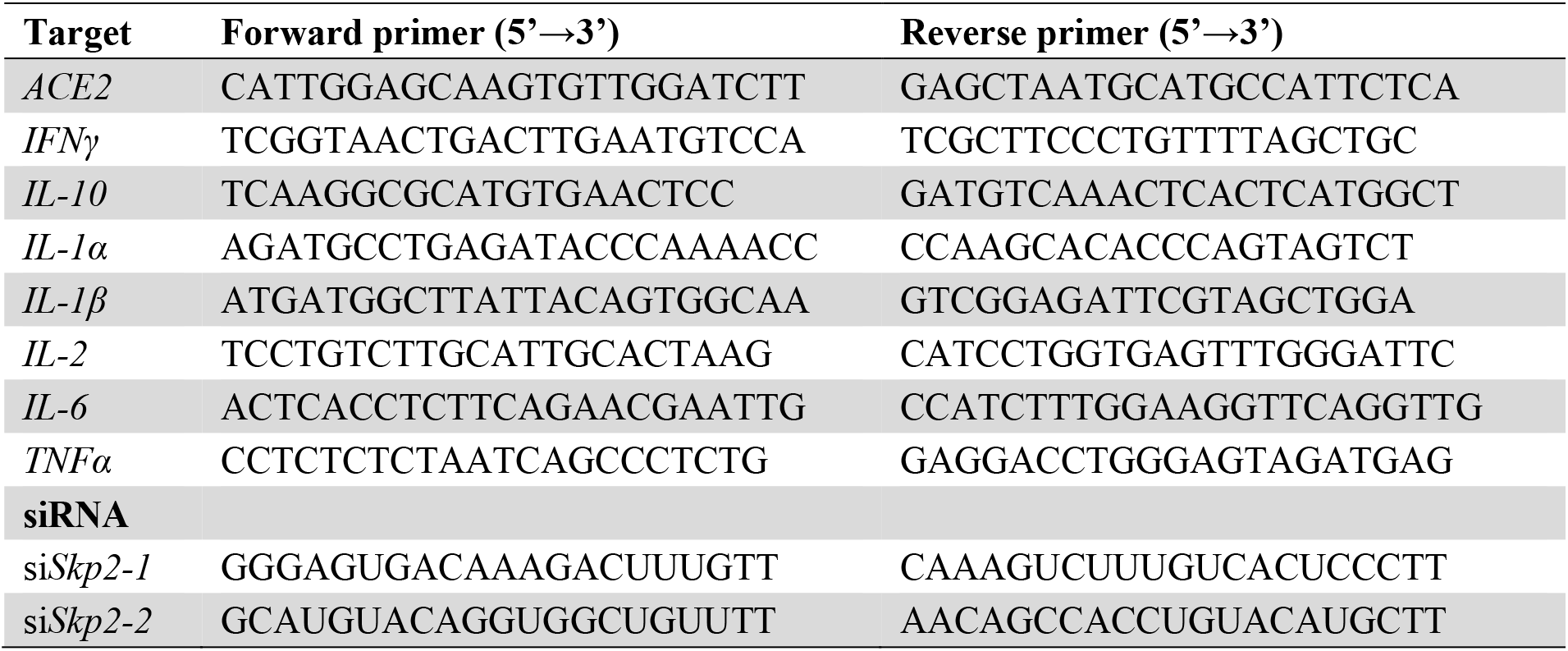
Primers and siRNAs used in the study.

**Supplementary Figure 1.**
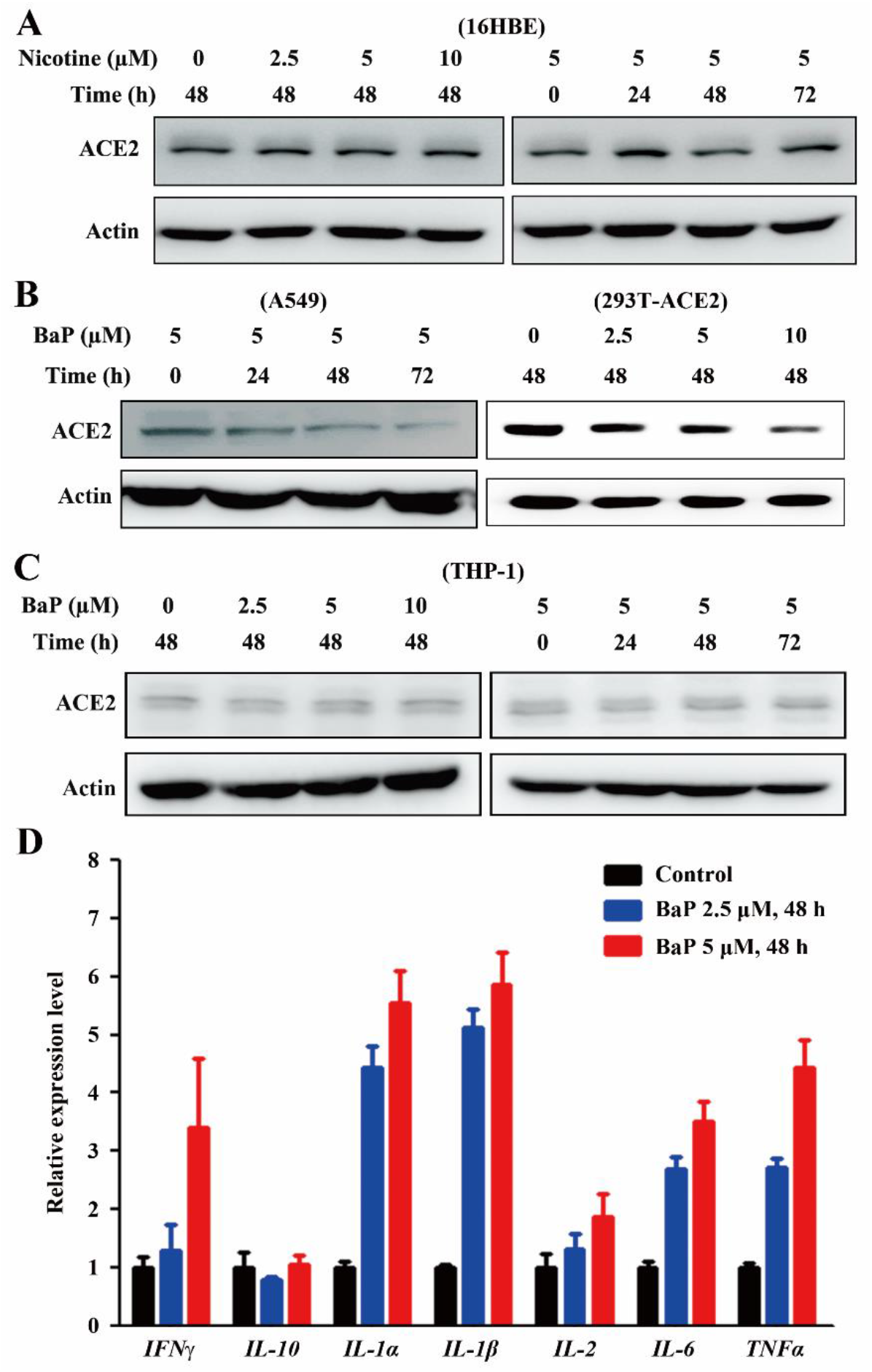
Effects of tobacco compounds on ACE2. (A - C) Indicated cells were treated with nicotine or BaP at indicated protocols, lysed, and subjected to Western blot. (D) THP-1 cells were treated with phorbol 12-myristate 13-acetate (PMA) at 100 ng/mL for 24 h, followed by BaP co-incubation for 48 h. The cells were lysed, RNA was extracted, and real-time PCR was conducted to evaluate the expression of indicated genes.

**Supplementary Figure 2.**
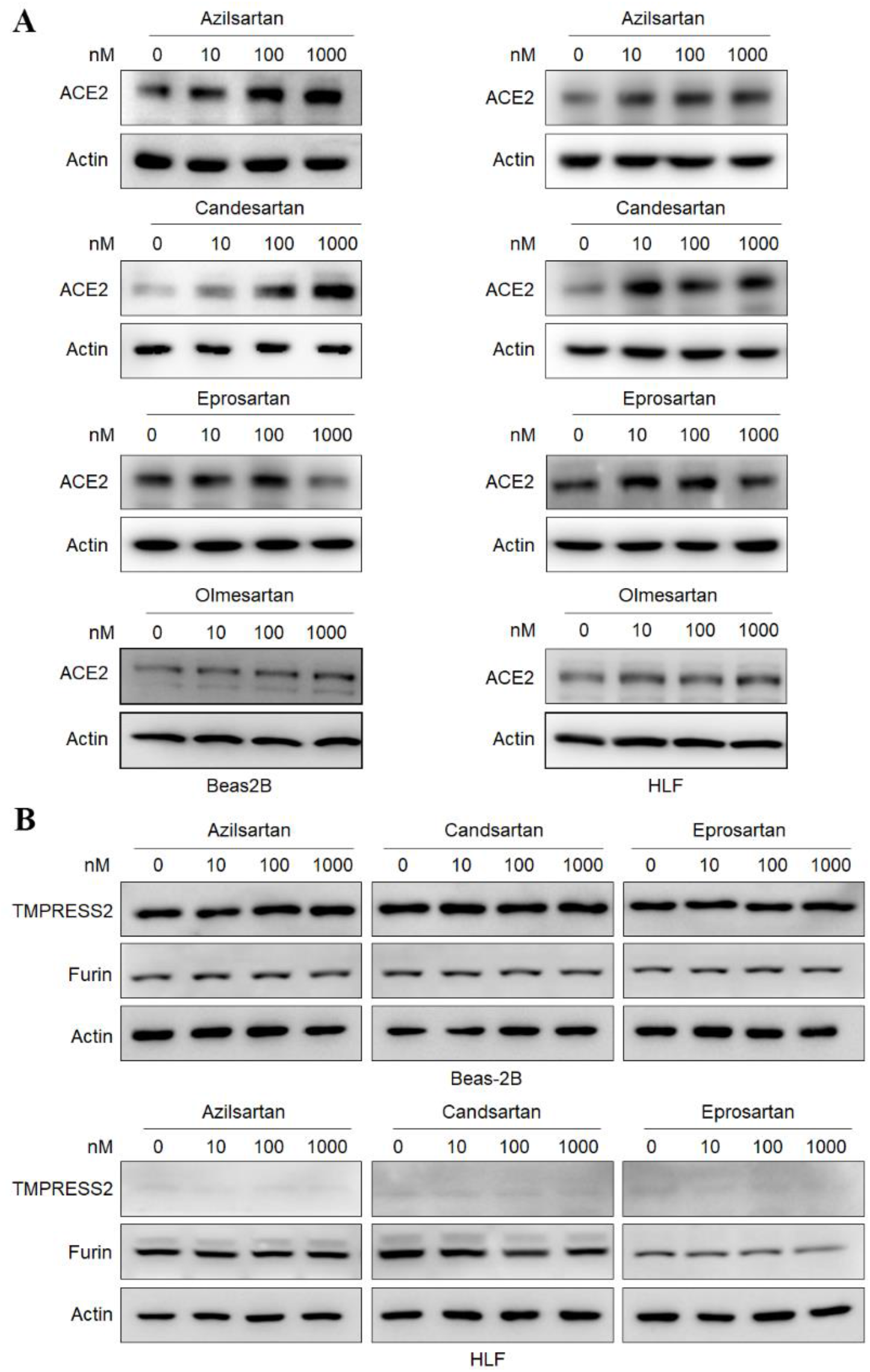
Effects of the angiotensin receptor blockers on the expression of ACE2 (A) and TMPRSS2 and Furin (B) in Beas-2B and HLF cells. The cells were treated indicated compounds at indicated concentrations for 48 h, lysed, and subjected to Western blot using indicated antibodies.

**Supplementary Figure 3.**
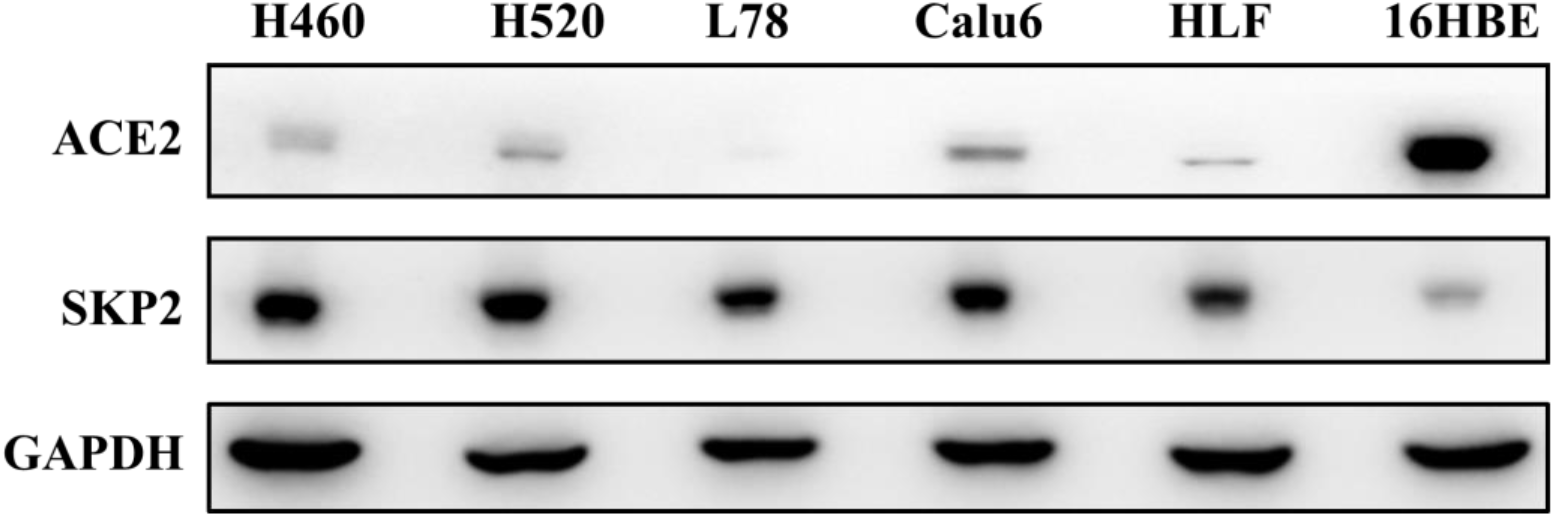
The expression ACE2 and Skp2 in 6 cell lines. H460 and H520, non-small cell lung cancer cell lines; L78, human lung squamous carcinoma cell line; Calu6, human pulmonary carcinoma cell line; HLF, human embryonic lung fibroblast cell line; 16HBE, human normal bronchial epithelial cell.

